# Revive-Flow: A Foundation Model for Blood DNAm Aging

**DOI:** 10.1101/2025.09.18.677241

**Authors:** Elon Litman, Tyler Myers, Vinayak Agarwal, Ekansh Mittal, Ashwin Gopinath, Timothy Kassis

## Abstract

Epigenetic clocks can predict biological age but cannot prescribe the interventions needed to reverse it. Here, we introduce REjuVenatIon Via Epigenetic Flow (Revive-Flow), a flow-matching model trained on a broad compendium of epigenetic blood studies to transport methylomes forward and backward in time. First, we learn the continuous vector field of aging as an Ordinary Differential Equation (ODE) within a stable, low-dimensional linear space. Then, the learned ODE’s dynamics are integrated backward in time to define a natural, biologically-plausible rejuvenation trajectory. This path serves as a guide for a convex optimization problem that identifies the minimal, targeted CpG-level perturbation required to rejuvenate a sample. On a completely unseen test set comprising over 800 individuals from the European Prospective Investigation into Cancer and Nutrition (EPIC-Italy) cohort, Revive achieves 0.4 years rejuvenated per commanded year (*R*^2^ ≈ 0.99), with a smooth sparsity-effect trade-off. Extensive validation confirms the effect preserves inferred cell-type composition and targets biologically plausible loci enriched in genomic regions and pathways central to aging biology.

## 1 Introduction

Aging leaves a measurable record in blood DNA methylation. Dozens of CpG sites shift in a predictable way with age, and learned “epigenetic clocks” can read this signal with striking accuracy (Horvath, 2013; Hannum et al., 2013; Levine et al., 2018; Lu et al., 2019; Myers et al., 2025). These clocks are valuable as biomarkers of health span and risk, but they are diagnostic rather than prescriptive. They tell us *how old* a methylome looks, not *how to move* it toward a younger state. Since epigenetic change is a recognized feature of organismal aging (López-Otín et al., 2013, 2023), a natural next step is to ask a concrete question:

What small, targeted set of CpG edits would most effectively rejuvenate a blood methylome?

We take a direct route to this question. Instead of generating many synthetic methylomes or relying on complex stochastic samplers, we model aging as a smooth flow through a low dimensional space and then solve a simple control problem on top of that flow. The idea is pragmatic; if we can learn a stable vector field that points in the direction methylomes tend to move as people age, we can integrate that field backward in time to define a plausible rejuvenation path. Then we can ask for the smallest change to the original CpGs that places a sample close to that path.

Our method, Revive, follows three steps. First, we learn a linear latent space with principal component analysis (PCA) for its stability, interpretability, and exact invertibility (Jolliffe, 2002). Second, within this space we learn a continuous time vector field of aging using an age calibrated flow objective that reduces training to plain regression, drawing on recent advances in flow matching and neural ordinary differential equations (Chen et al., 2018; Lipman et al., 2022; Tong et al., 2023; Liu et al., 2022). Third, we turn the rejuvenation goal into a convex optimization that trades fidelity to the natural backward trajectory against sparsity of CpG edits and protection of low variance loci, and we solve it efficiently with proximal updates via Alternating Direction Method of Multipliers (ADMM) (Boyd et al., 2011; Parikh and Boyd, 2014).

We build a large blood compendium by filtering MethAgingDB (Li et al., 2025) to human blood studies with modern arrays and at least four hundred thousand CpGs, intersect CpGs across studies, and standardize within each study to reduce unwanted technical variation (Aryee et al., 2014; Fortin et al., 2017; Leek et al., 2010). One complete cohort, EPIC-Italy (GSE51032), is held out from start to finish. It does not influence preprocessing, principal components, the learned dynamics, or hyperparameters, and serves as a strict unseen test set. Because bulk blood methylation reflects both within cell type aging and changing leukocyte mixtures, we also verify that our edits do not simply reshuffle inferred cell type proportions using a standard reference based deconvolution (Houseman et al., 2012).

In summary, Revive turns epigenetic aging from a readout into a controllable process. A simple linear embedding, a learned continuous flow tied to calendar age, and a small convex controller together yield targeted CpG interventions that are dose controllable, sparse, and biologically plausible. The result is a practical framework for in silico design of epigenetic rejuvenation strategies in blood that complements existing clocks rather than replacing them.

## 2 Method

Our method, Revive, consists of three main stages, outlined in Figure 1: (1) learning a stable, low-dimensional manifold of epigenetic aging from harmonized DNAm data; (2) modeling the dynamics of aging within this manifold as a continuous-time ODE trained with a flow-matching objective; and (3) solving a convex optimization problem to translate a natural rejuvenation trajectory into a sparse, targeted intervention. We detail each stage in the following sections.

**Figure 1:**
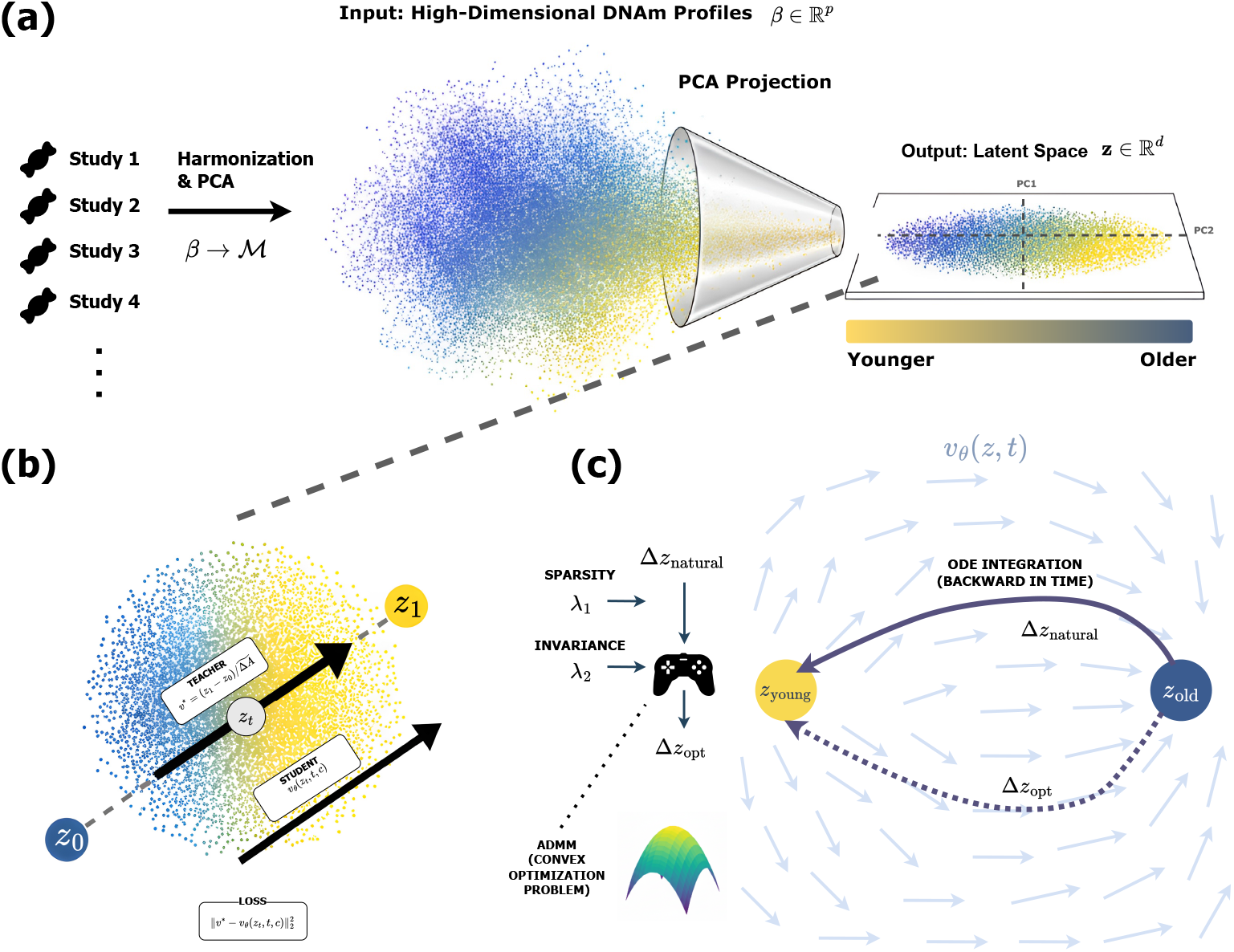
The Revive-Flow Pipeline. **(a) Data Harmonization and Manifold Learning:** Raw blood DNAm profiles (*β*) from multiple studies are transformed to M-values, standardized per-study, and projected into a stable, low-dimensional latent space (***z*** ∈ ℐ^*d*^) using Principal Component Analysis (PCA). **(b) Learning Aging Dynamics with Flow Matching:** We train a neural ODE model, *v*_*θ*_, to learn the vector field of aging. The model is trained via a simple regression objective to match the time-calibrated velocity between pairs of samples (***z***_0_, ***z***_1_) interpolated at a random time *t*. **(c) Sparse Intervention Design:** For a given sample ***z***_old_, we first define a natural rejuvenation trajectory by integrating the learned ODE backward in time to a target age, yielding Δ***z***_natural_. This vector guides a convex optimization problem, solved via ADMM, that finds an optimal perturbation Δ***z***_opt_ balancing fidelity to the natural path with sparsity (λ_1_) and biological invariance (λ_2_). This optimal latent change is then decoded back to a minimal set of targeted CpG edits (Δ*β*^∗^).

### 2.1 Dataset & Preprocessing

All preprocessing, PCA, model training, and validation are conducted for blood tissue only. We enumerate all available blood studies (datasets), compute the CpG intersection across them, and evaluate our model using a rigorous train/test split. We pool all available blood datasets for training, with the exception of one large, independent cohort: GSE51032 from the EPIC-Italy study, which is held out entirely from all preprocessing, model fitting, and hyperparameter tuning to serve as a strict, unseen test set.

#### Data Structures and Notation

Let 𝒟 = {*D*_1_, …, *D*_|*D*|_ } be a collection of independent blood DNAm datasets. The total number of subjects across all datasets is *N*, and the initial number of CpG features is *p*_0_. For each subject *n* ∈ {1, …, *N* }, we assume the availability of the following data tuple: *β*_*n*_ ∈ [0, 1]^*p*0^, a vector of CpG methylation levels (beta-values); *A*_*n*_ ∈ ℝ^+^, the chronological age; and *S*_*n*_ ∈ {Male, Female}, the annotated sex. *D*_*n*_ ∈ 𝒟 is an indicator for the dataset of origin.

#### CpG Harmonization

To ensure a consistent feature space, we define the CpG set ℐ as the intersection of all CpG sites present across all blood datasets in our compendium. All methylation profiles *β*_*n*_ are restricted to this set ℐ, resulting in new vectors of dimension *p* = |ℐ|, which we continue to denote as *β*_*n*_ for simplicity.

#### Transformation and Per-Dataset Standardization

To stabilize variance and mitigate technical batch effects arising from different studies, we employ a two-step process. First, beta-values are converted to M-values using the logit transformation, which maps the bounded [0, 1] interval to the real line ℝ.

##### Definition 2.1: M-value Transformation

For a methylation profile *β*_*n*_, the corresponding M-value profile ***M***_*n*_ ∈ ℝ^*p*^ is given by:

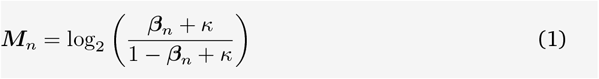

where *κ* = 10^−6^ is a small offset added for numerical stability.

##### Definition 2.2: Inverse M-value Transformation

The mathematically precise inverse transformation from an M-value profile ***M***_*n*_ back to beta-values is:

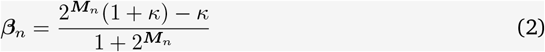

This formulation ensures that the forward and inverse transforms are consistent.

Second, to harmonize data across different studies, we standardize the M-values within each dataset. For each dataset *d* ∈ 𝒟, we compute the feature-wise mean ***µ***_*d*_ and standard deviation ***σ***_*d*_ over all samples belonging to that dataset. Each sample ***M***_*n*_ from dataset *d* is then standardized:

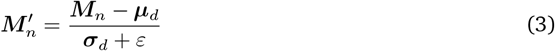

where *ℰ*= 10^−8^ is a small constant to prevent division by zero. For a held-out test study *d*^∗^, the moments 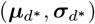 are computed *unsupervised over all samples of that held-out study* and reused for all of its test-time preprocessing.

### 2.2 Learning the Dynamics of Aging

The first phase of our methodology is to learn a robust, continuous-time model of the blood aging process. This is accomplished by identifying a low-dimensional manifold and then learning a vector field on that manifold.

#### Stable Manifold Estimation via Principal Component Analysis

We project the high-dimensional standardized M-values into a low-dimensional and computationally tractable space using Principal Component Analysis (PCA). We explicitly choose PCA over non-linear alternatives for its determinism, geometric simplicity, and efficacy as a denoising filter. The PCA model is fit on the pooled, standardized training data from all *non-held-out* blood studies; the resulting mean and loadings are then fixed and reused at test time. Let 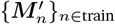 be the collection of all standardized profiles from the designated training datasets. We first compute their global mean,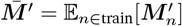. PCA is then performed on the centered data matrix 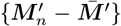 to find the loading matrix *W* ∈ ℝ^*p*×*d*^, whose columns are the top *d* orthonormal eigenvectors of the covariance matrix. The latent dimension *d* ≪ *p* is a hyperparameter set to 1024, explaining 67% of the variance.

##### Definition 2.3: Latent Space Projection and Decoding

The latent representation ***z***_*n*_ ∈ ℝ^*d*^ for any standardized sample 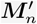 is a linear projection:

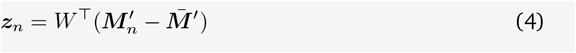

The inverse projection, or decoding, from the latent space back to the standardized M-value space is:

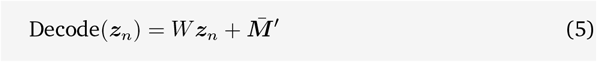

#### Modeling Aging as a Continuous-Time ODE

We model the process of aging as a continuous trajectory within the PCA latent space. This trajectory is governed by a learned vector field that is conditioned on the sample’s current state, its chronological age, and its biological context. Let ***z***(*t*) be the state at a continuous time (age) parameter *t*. Its evolution is described by the Ordinary Differential Equation (ODE):

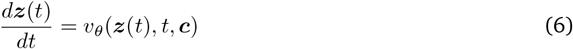

Here, *v*_*θ*_ : ℝ^*d*^ × ℝ^+^ × ℝ^|***c***|^ → ℝ^*d*^ is a neural network with parameters *θ* that defines the vector field. The condition vector ***c*** provides biological context (sex), and the model is explicitly conditioned on the absolute age *t*.

#### Learning with an Age-Calibrated Flow Objective

We learn the parameters *θ* of the vector field *v*_*θ*_ using an objective inspired by conditional flow matching. This approach is powerful in its simplicity and stability. It posits that the path between any two observed data points can be approximated by a straight line in our linear PCA space, and the model’s task is to learn the time-calibrated velocity along this path.

We train with pairs of samples (***x***_0_, ***x***_1_) from the blood training set, where ***x***_*i*_ = (***z***_*i*_, *A*_*i*_, *S*_*i*_). To ensure the conditioning is valid, we only sample pairs that share the same sex, *S*_0_ = *S*_1_. To expose the model to both aging and rejuvenation directions, we apply a random bidirectional flip: with probability 1/2, we swap the order of the pair. Let the post-swap pair be (***z***_0_, *A*_0_) and (***z***_1_, *A*_1_). We define a point on the linear interpolation path as ***z***_*s*_ = (1− s)***z***_0_ + *s****z***_1_ for *s* ∈ [0, 1], corresponding to the interpolated age *t*_*s*_ = (1−*s*)*A*_0_ + *sA*_1_. The crucial insight is to define a teacher velocity that is calibrated to chronological time. The velocity with respect to the path parameter s is ***z***_1_ − ***z***_0_. To prevent exploding targets for very small age gaps, we floor the signed age difference:

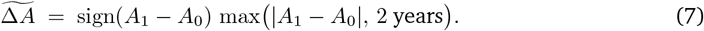

The time-calibrated teacher velocity is then

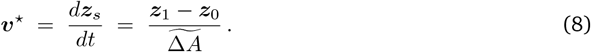

This gives a physically meaningful target with units of “change in latent space per year”.

##### Definition 2.4: Age-Calibrated Flow Loss

The learning objective is to minimize the expected squared Euclidean distance between the model’s predicted vector field and the time-calibrated teacher velocity. The teacher velocity is defined using a stabilized age denominator, 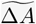, to prevent numerical instability with small age gaps. The loss is:

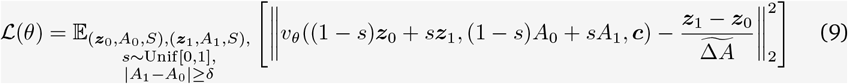

where the condition vector ***c*** contains a one-hot encoding of the sex *S, δ* is a minimum age gap threshold (e.g., *δ* = 5), and *A*_min_gap_ is the floor (e.g., 2 years). This objective is a simple mean squared error regression problem, guaranteeing stable training.

#### Model Architecture

We parameterize the vector field network *v*_*θ*_ with a Transformer encoder to model dependencies between the principal components (PCs) of the methylome. The latent vector ***z*** ∈ ℝ^*d*^ is treated as a sequence of *d* tokens, where each token is the scalar score of a single PC that is linearly projected into a high-dimensional embedding. The model is conditioned on chronological age *t*, encoded as sinusoidal Fourier features, and sex, represented as a one-hot vector. This combined context generates a scale (*γ*) and shift (*β*) for each token embedding via a Feature-wise Linear Modulation (FiLM) layer. These modulated embeddings are then processed by the Transformer’s self-attention layers. A final linear head reduces each output embedding back to a single scalar, forming the resultant vector field *v*_*θ*_ ∈ ℝ^*d*^. The full procedure for data preprocessing, PCA, and training the dynamics model is summarized in Algorithm 1.

### 2.3 Identifying Sparse Interventions

With the trained dynamics model *v*_*θ*_, we now address the core goal: identifying minimal, targeted interventions for rejuvenation.

#### The Rejuvenation Trajectory via ODE Integration

The trained model *v*_*θ*_ defines a continuous vector field for aging. The most natural, biologically congruent path for rejuvenation is therefore the trajectory obtained by following this field backward in time.

##### Definition 2.5: Natural Rejuvenation Trajectory

For an aged sample with latent representation ***z***_old_ at age *A*_old_, and a desired target age *A*_young_ < *A*_old_, the naturally rejuvenated state ***z***_young_ is the solution to the ODE initial value problem integrated backward in time:

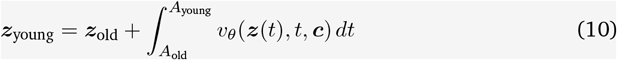

We compute this integral with a fixed-step fourth-order Runge–Kutta (RK4) scheme using 300 uniform steps.^*a*^ We also clamp the target age to a minimum feasible value *A*_min_ = 5 years and integrate to *A*_young_ ← max(*A*_young_, *A*_min_) for numerical/biological safety. The total displacement from this integration defines the natural rejuvenation vector, Δ***z***_natural_ = ***z***_young_ − ***z***_old_. This vector represents the ideal, non-linear change in the latent space to achieve rejuvenation according to the learned dynamics.

#### Solving the Sparse Control Problem in *M*-Space

While applying Δ***z***_natural_ directly would produce a biologically plausible rejuvenated profile, it would likely correspond to dense, small changes across thousands of CpGs, making it therapeutically impractical. We therefore seek an optimal perturbation, Δ***z***_opt_, that is close to the natural vector but corresponds to a sparse set of edits in the original CpG space. This is formulated as a convex optimization problem.

##### Definition 2.6: Sparse Rejuvenation Trajectory

Let 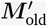 be the standardized M-value profile of the aged sample, which originated from study *D*_old_ with standard deviations 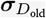. We solve for the optimal latent perturbation Δ***z*** by minimizing a composite objective function:

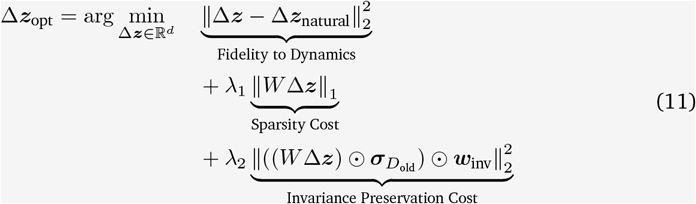

The three terms are: a fidelity term anchoring the solution to the natural rejuvenation dynamics; a sparsity-inducing ℓ_1_ term on the CpG-aligned standardized change ***q*** = *W* Δ***z*** in *M*^′^-space; and an invariance preservation term. Sparsity is therefore imposed in standardized *M*^′^ space; reported CpG edit counts in *β*-space use a threshold of |Δ*β*| > 10^−3^ after the inverse link. This third term penalizes the squared change in the *unstandardized* M-value space, 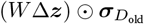, weighted to protect CpG sites with low population variance. The invariance weights are pre-computed from the variance of the unstandardized training M-values, per CpG *k*:

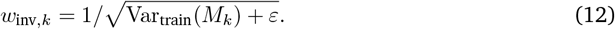

This problem is convex and is particularly well-suited for the Alternating Direction Method of Multipliers (ADMM). We introduce an auxiliary variable ***q*** = *W* Δ***z*** to decouple the terms. The problem becomes:

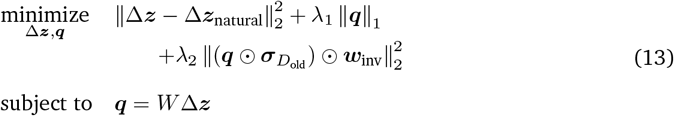

The ADMM algorithm iteratively solves for Δ***z*** and ***q*** and then updates a dual variable. The update for ***q*** involves a proximal operator for an elastic-net-like penalty, which has a closed-form solution involving soft-thresholding. This approach is superior to generic gradient-based methods for non-smooth objectives. For enhanced biological safety, box constraints of the form |Δ*β*_*k*_| ≤ Δ_max_ or masks for functionally protected loci (e.g., imprinted genes) can also be incorporated into the optimization.

#### Generating the Final Intervention in *β*-Space

After solving for Δ***z***_opt_, the final rejuvenated latent state ***z***_opt_ = ***z***_old_ + Δ***z***_opt_ is decoded back to the beta-value space by reversing all preprocessing steps, using the moments 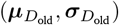 from the sample’s original dataset *D*_old_. We preserve each sample’s non-PCA structure by adding back its PCA residual in standardized space. Let

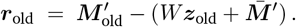

Then the decoding is

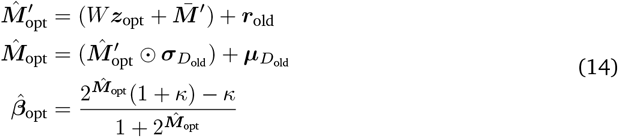

The final intervention is the sparse vector of beta-value changes, 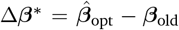. This entire three-step process for generating a targeted intervention is detailed in Algorithm 2.

## 3 Results

We evaluated Revive on the EPIC-Italy GSE51032 cohort, a dataset strictly held out from all model training and hyperparameter selection. To ensure an unbiased assessment, we employed an internal judge: a Ridge regressor trained on the standardized M-values (***M***^*′*^) of the complete training set. Its ℓ_2_ regularization and linear nature provide a robust and conservative measure of age modification. All reported realized ΔAge values are the difference in this judge’s age predictions before and after an intervention.

### 3.1 Revive Achieves Linearly Controllable Epigenetic Rejuvenation

#### Efficacy: Commanded vs. Realized Rejuvenation

For each held-out sample, we command rejuvenations of Δ*a* ∈ {2, 5, 7, 10} via the ADMM controller, predict age with a Ridge judge trained on **M**^*′*^, and for each (λ_1_, Δ*a*) report the mean realized ΔAge with its 95% t-interval. Across studies, the aggregate slope is 0.291 ± 0.148 with study-wise *R*^2^ = 0.910 ± 0.116; in GSE51032 the slope is 0.396 with *R*^2^ = 0.985 at λ_1_ = 0. In Figure 2, the dose–response is linear and the slope (years rejuvenated per commanded year) decreases with higher sparsity (larger λ_1_), confirming the expected trade-off.

**Figure 2:**
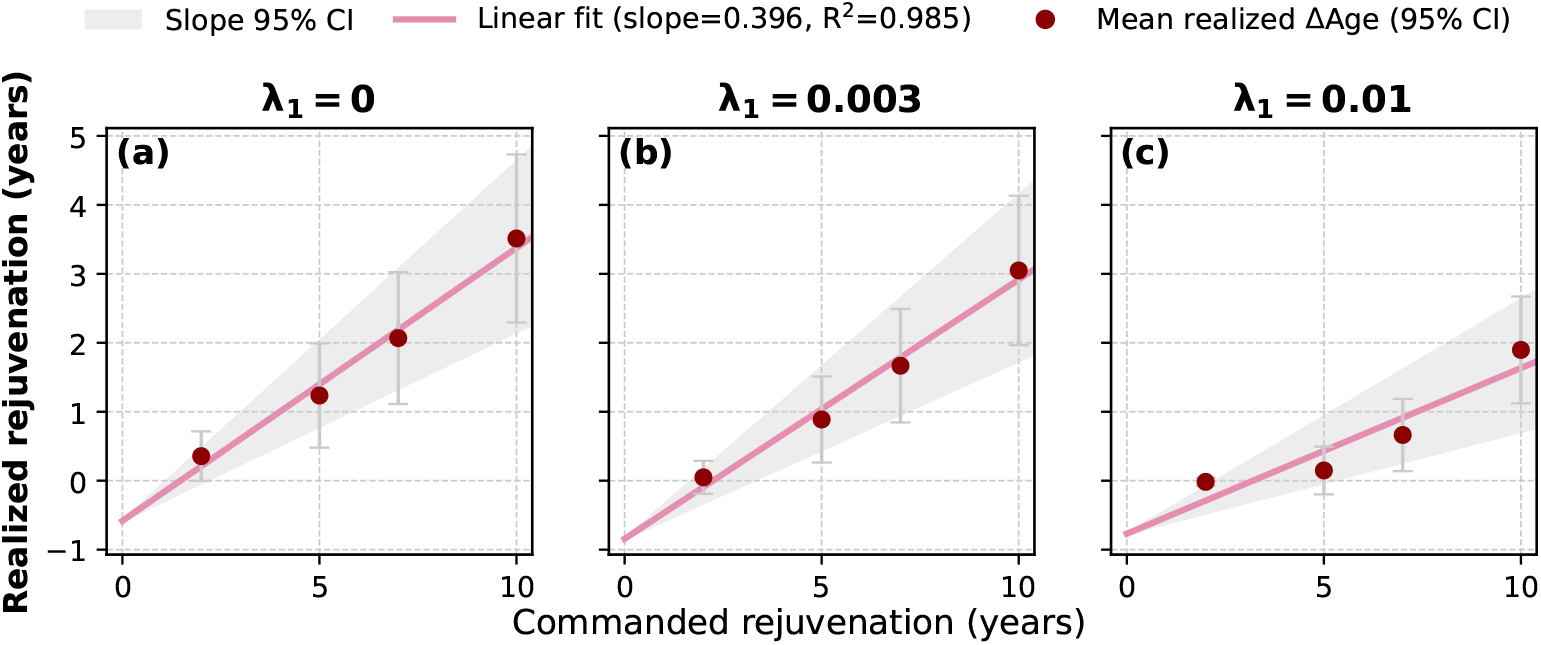
Efficacy on the held-out study: commanded vs. realized rejuvenation for three *λ*_1_ settings. Panels: **(a)** low sparsity penalty (λ_1_ = 0), **(b)** moderate (λ_1_ = 0.003), **(c)** high (λ_1_ = 0.01). Each point is the cohort mean over subjects for one commanded Δa. Lines show least-squares fits; shaded bands are 95% bootstrap CIs for the slope.

We regress realized on commanded ΔAge across Δ*a* (per λ_1_), reporting slope and *R*^2^. Table 2 summarizes the held-out study and across-study mean. Per–(λ_1_, Δ*a*) results are reported in Table 1.

**Table 1:** Mean realized ΔAge (years) on EPIC-Italy at each commanded Δ*a* (95% t–intervals).

| $\lambda_1$ | $\Delta a = 2 \text{ y}$ | $\Delta a = 5 \text{ y}$ | $\Delta a = 7 \text{ y}$ | $\Delta a = 10 \text{ y}$ |
| --- | --- | --- | --- | --- |
| 0.000 | 0.356 [-0.003, 0.715] | 1.236 [0.480, 1.992] | 2.068 [1.111, 3.025] | 3.513 [2.294, 4.731] |
| 0.001 | 0.256 [-0.061, 0.574] | 1.123 [0.411, 1.834] | 1.937 [1.025, 2.849] | 3.360 [2.187, 4.534] |
| 0.003 | 0.048 [-0.192, 0.288] | 0.888 [0.264, 1.512] | 1.668 [0.844, 2.491] | 3.049 [1.965, 4.133] |
| 0.010 | -0.017 [-0.075, 0.042] | 0.149 [-0.198, 0.495] | 0.663 [0.139, 1.187] | 1.897 [1.123, 2.671] |
| 0.020 | -0.00086 [-0.00226, 0.00054] | 0.00314 [-0.0957, 0.102] | 0.0556 [-0.143, 0.254] | 0.428 [0.0436, 0.813] |

**Table 2:** Commanded realized linearity on GSE51032. Slope is years realized per commanded year (95% CI); *R*^2^ from the same fit.

| $\lambda_1$ | Slope [95% CI] | $R^2$ |
| --- | --- | --- |
| 0.000 | 0.396 [0.274, 0.521] | 0.985 |
| 0.001 | 0.389 [0.270, 0.513] | 0.985 |
| 0.003 | 0.376 [0.258, 0.498] | 0.985 |
| 0.010 | 0.240 [0.148, 0.339] | 0.873 |
| 0.020 | 0.052 [0.0039, 0.101] | 0.722 |

**Table 3:**
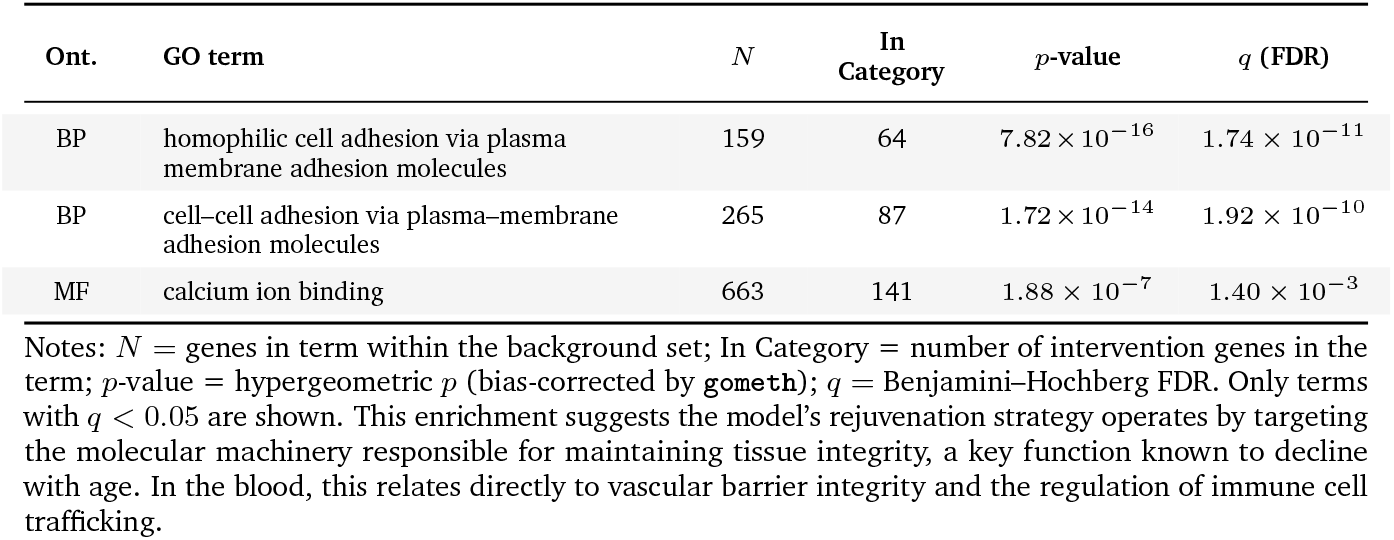
GO enrichment for Intervention CpG genes (gometh).

| Ont. | GO term | $N$ | In Category | $p$-value | $q$ (FDR) |
| --- | --- | --- | --- | --- | --- |
| BP | homophilic cell adhesion via plasma membrane adhesion molecules | 159 | 64 | $7.82 \times 10^{-16}$ | $1.74 \times 10^{-11}$ |
| BP | cell–cell adhesion via plasma–membrane adhesion molecules | 265 | 87 | $1.72 \times 10^{-14}$ | $1.92 \times 10^{-10}$ |
| MF | calcium ion binding | 663 | 141 | $1.88 \times 10^{-7}$ | $1.40 \times 10^{-3}$ |

Notes: $N$ = genes in term within the background set; In Category = number of intervention genes in the term; $p$ -value = hypergeometric $p$ (bias-corrected by *gometh*); $q$ = Benjamini–Hochberg FDR. Only terms with $q < 0.05$ are shown. This enrichment suggests the model’s rejuvenation strategy operates by targeting the molecular machinery responsible for maintaining tissue integrity, a key function known to decline with age. In the blood, this relates directly to vascular barrier integrity and the regulation of immune cell trafficking.

#### Controllability: Sparsity–Effect Trade-off

We quantify controllability by the trade-off between realized rejuvenation and the number of edited CpGs (counting |Δ*β*| > 10^−3^). We aggregate, for each Δ*a*, the cohort mean edits and the mean realized ΔAge across λ_1_. Figure 3 visualizes this frontier: as λ_1_ decreases (rightward), realized ΔAge rises monotonically with the number of edited CpGs. This reflects a predictable, dose-like trade-off: denser edits yield larger rejuvenation, with a mild saturation at the highest edit counts. In practice, λ_1_ provides a single, transparent knob to choose an edit budget that matches a desired rejuvenation level.

**Figure 3:**
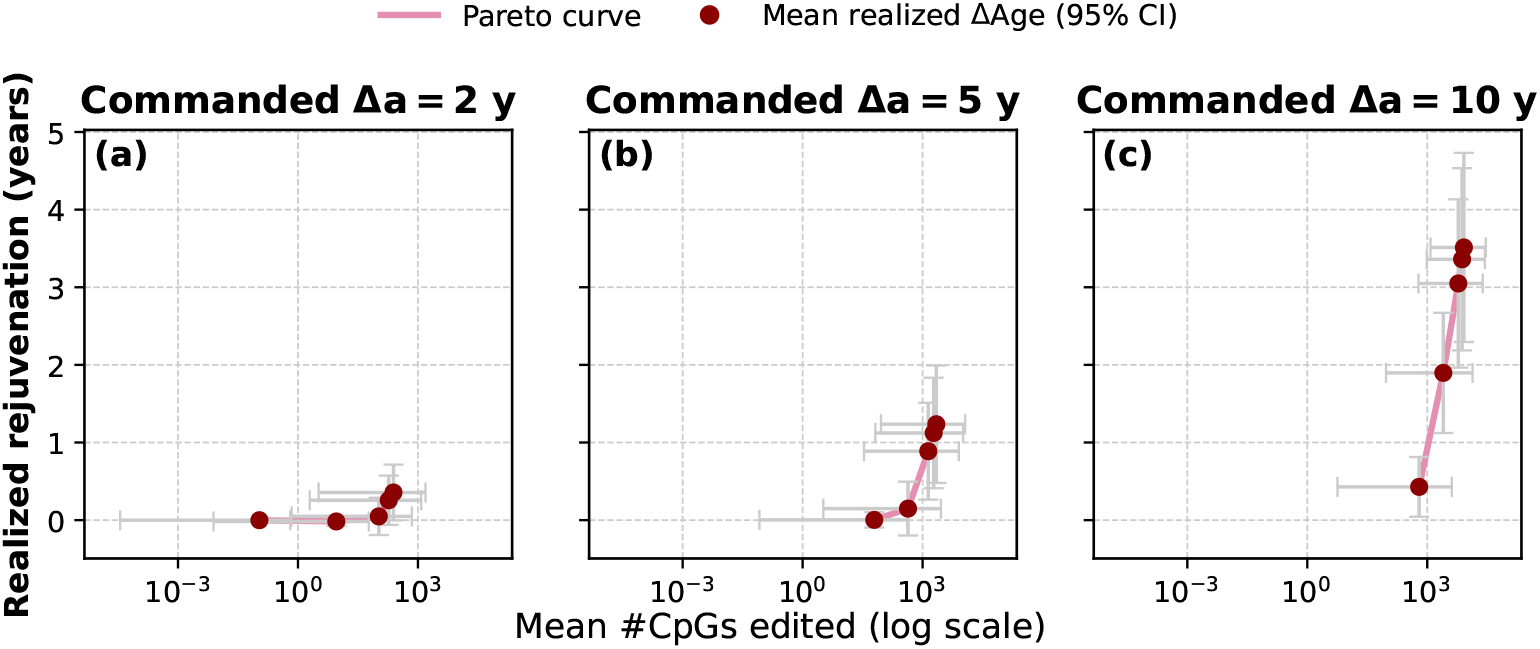
Pareto trade-off between sparsity and rejuvenation (EPIC-Italy hold-out). For commanded rejuvenation Δ*a* ∈ {2, 5, 7, 10} years, realized ΔAge (y-axis) increases monotonically with the mean number of edited CpGs (x-axis, log_10_ scale). *Points* show cohort means with 95% CIs for realized ΔAge; *horizontal error bars* show the SD of #CpGs edited across subjects. The *pink line* is the *Pareto envelope across sparsity settings* (λ_1_): moving left corresponds to higher λ_1_ (sparser), moving right to lower λ_1_ (denser). Diminishing returns are evident beyond ~ 10^3^ − 10^4^ edits. Edits are counted in β-space with threshold |Δβ| > 10^−3^ (section 2). Common y-limits are used across panels for comparability; small effects at Δ*a* = 2 years therefore appear visually compressed but remain positive.

### 3.2 Ablations and Validation Confirm Model Integrity

We subject Revive to targeted ablations and experiments to rule out artifactual gains, leakage, or overfitting and to verify that the controller’s effect is biologically and statistically valid. First, a shuffled-age negative control destroys the chronological-age signal and shows that commanded rejuvenation does not spuriously appear; realized changes are non-positive with negative realized-vs-commanded slopes, indicating the pipeline does not fabricate improvement. Second, we examine cell-type composition and find minimal total-variation shifts and small mean absolute changes per cell type, showing that bulk rebalancing cannot explain the observed rejuvenation. Across these checks (with bootstrap CIs and sensitivity to sparsity), the evidence is consistent: the model’s edits act on age-linked methylation rather than on confounds, supporting the integrity of our main results.

#### 3.2.1 Negative Control

We repeated the entire pipeline after *randomly permuting donor ages* while keeping all other steps identical (EPIC-Italy hold-out). Under this null, commanded rejuvenation should not translate into realized rejuvenation; the relationship between realized and commanded changes should be near zero or negative.

The shuffled-age control behaves accordingly. In Figure 4a, positive commands yield non-positive realized changes (mean ± 95% CI) across Δ*a* ∈ {2, 5, 7, 10} years. Figure 4b shows realized-vs-commanded slopes that are consistently negative, approaching zero only under the strongest penalization. This indicates the procedure does not fabricate rejuvenation when chronological age is destroyed, and supports the main result that Revive *reverses* aging rather than exploiting artifacts or leakage.

**Figure 4:**
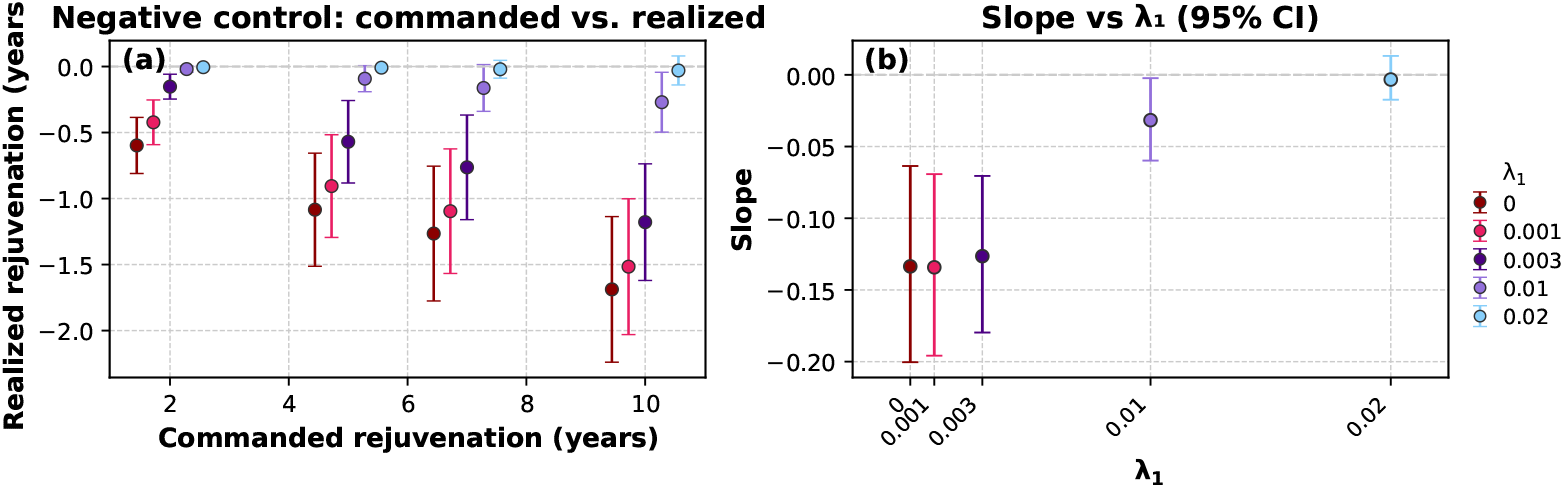
Negative control on EPIC-Italy (shuffled donor ages). **(a)** Commanded vs. realized rejuvenation (mean ± 95% CI) for Δ*a* ∈ {2, 5, 7, 10} years across λ_1_ values. Realized changes move in the wrong direction (aging). **(b)** Slope of realized versus commanded as a function of λ_1_ with 95% CIs; slopes are negative and only approach zero at the strongest penalty. When chronological age is randomized, the pipeline does not produce spurious rejuvenation, supporting the validity of the positive effects observed in the unshuffled runs.

#### 3.2.2 Cell-type Composition is Preserved by the Controller

Edits that merely reshuffle inferred blood cell-type proportions could artifactually change age readouts. We therefore test whether our controller alters *composition* or primarily the age-related methylation signature within cell types.

#### Setup and metrics

For each held-out cohort and operating point (commanded Δ*a* ∈ {2, 5, 7, 10} years; three values of λ_1_ spanning sparse→dense solutions), we estimate cell-type fractions on the *original* and *edited* methylomes using the same reference-based deconvolution (Houseman method). Let 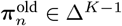 and 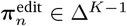 denote the inferred *K*-part compositions for sample *n*. For each cell type *k*, we report the mean absolute change (MAC),

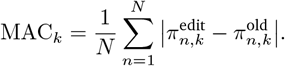

Per sample, we summarize the overall composition shift by total variation (TV),

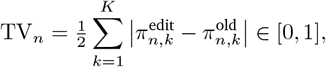

which equals 0 when compositions are unchanged. Figure 5 presents three complementary checks. First, Figure 5a shows mean absolute change (MAC) per cell type with 95% CIs, which are uniformly small, indicating no systematic rebalancing of any lineage. Second, Figure 5b displays per-sample Δ*π* values that are narrowly centered around zero across samples and cell types. Third, Figure 5c summarizes the overall shift with total variation (TV): in EPIC-Italy at Δ*a* = 10 years and λ_1_ = 0.003, TV is tightly concentrated near zero. Together, these show that the controller leaves inferred composition essentially unchanged, so the observed rejuvenation is explained by within-cell-type methylation changes rather than bulk cell-type mixing.

**Figure 5:**
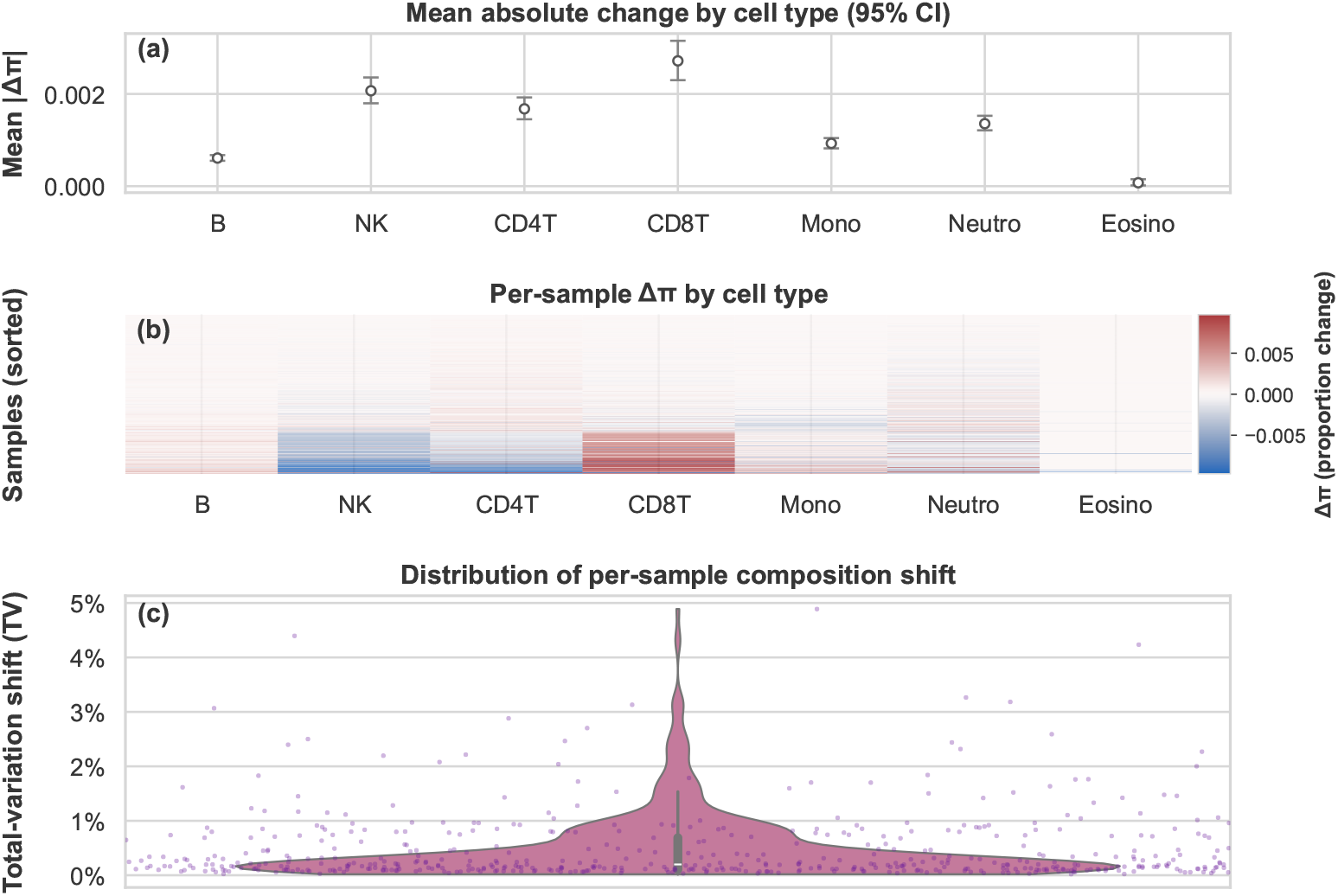
Cell-type stability under edits (EPIC-Italy hold-out; Δ*a* = 10 years, *λ*_1_ = 0.003). **(a)** Mean absolute change by cell type (MAC; mean ± 95% CI) is uniformly small. **(b)** Heatmap of per-sample Δ*π* (rows denote samples sorted by realized ΔAge; columns denote cell types) **(c)** Distribution of per-sample total-variation shift (TV) shows minimal composition change (median 0.2%, 95^th^ percentile 1.7%). The small TV magnitudes indicate that the observed rejuvenation is not explained by bulk cell-type rebalancing.

### 3.3 Biological Plausibility: Intervention Targets are Enriched for Aging Hallmarks

Lastly, we demonstrate the biological plausibility of identified intervention targets. To do this, we ask where, in the genome, the intervention-selected CpGs tend to fall. Using the Illumina EPIC manifest we annotated every CpG in the background set by (i) relation to CpG islands (Island, Shore, Shelf, Open Sea) and (ii) gene feature/TSS group (TSS200, TSS1500, 1st Exon, 5’UTR, 3’UTR, Body). A global *χ*^2^ test showed that intervention status depends strongly on genomic context, and category–wise Fisher tests (BH–FDR) indicated the direction of effects. Intervention CpGs are enriched in *Islands and Shores* and depleted in *Shelves/Open Sea* (Figure 6a); by gene feature they are enriched in the *Gene Body* and *1st Exon*, approximately neutral in *5’UTR*, and depleted near the promoter/TSS regions (*TSS200/TSS1500*) and in *3’UTR* (Figure 6b). The intersection of Figure 6a and Figure 6b shows a preference for gene-associated CpGs, including islanded sites within first exons and bodies, while largely avoiding promoter-proximal CpG-dense regions.

**Figure 6:**
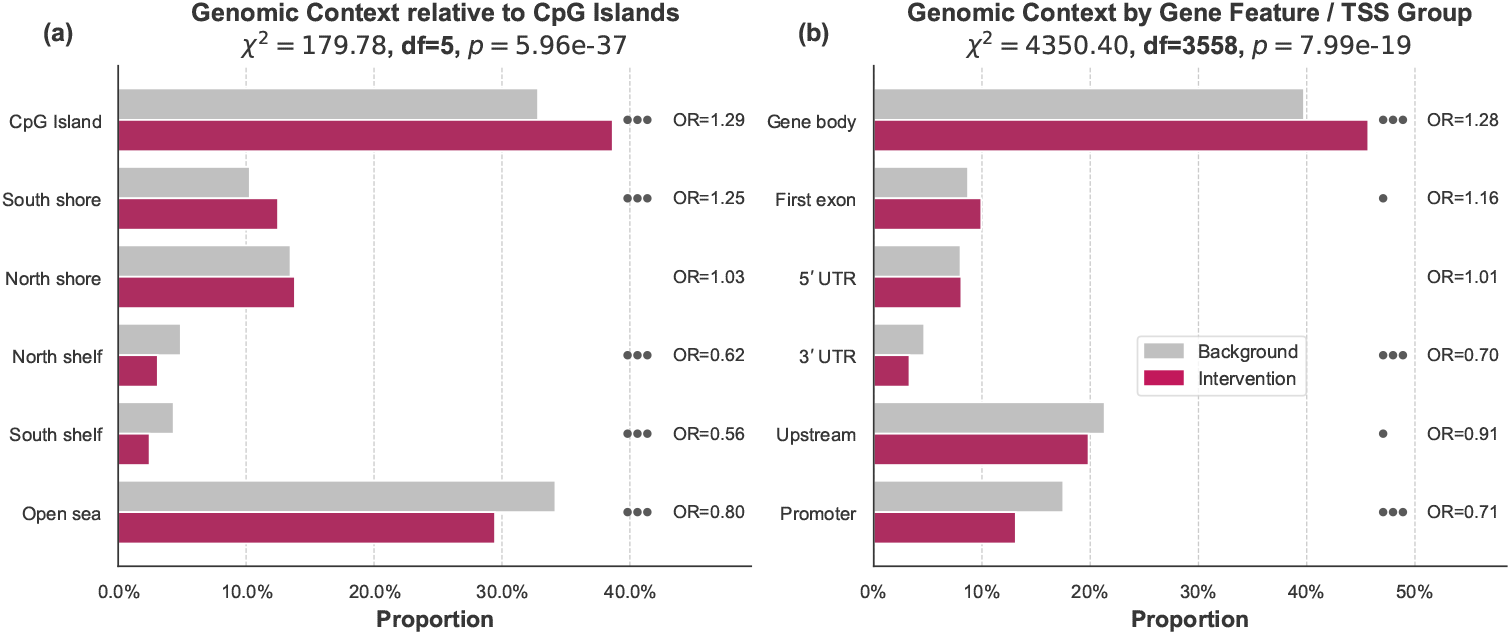
Genomic context enrichment (EPIC–Italy hold-out; Δ*a* = 10 years, *λ*_1_ = 0.003). **(a)** Context relative to CpG islands. **(b)** Context by gene feature / TSS group. Panel titles report global *χ*^2^ statistics with exact *p*-values computed from the contingency tables. “OR” labels denote the odds ratio for each category, computed from the 2 × 2 table of Intervention vs. Background; it is the ratio of the category’s proportion in the intervention set to that in the background (OR > 1 enrichment, OR < 1 depletion). Dots indicate Benjamini-Hochberg False Discovery Rate from Fisher’s exact tests per category (• *q* < 0.05, • • *q* < 0.01, • • • *q* < 0.001).

This pattern is expected. Promoter CpG islands are typically constitutively unmethylated and tightly constrained; altering them risks broad transcriptional effects. In contrast, CpGs in shores and within gene bodies/first exons exhibit greater dynamic range across development, aging, and cell state, making them more responsive, and safer, targets for modest methylation edits. The model therefore concentrates on contexts where small changes are informative and controllable, rather than on core promoters where methylation is less variable.

To understand the functional consequence of these edits, we mapped the Intervention CpG Set to their associated genes and performed a pathway enrichment analysis using the `gometh` method in **R**, which corrects for the non-uniform distribution of CpGs per gene. The analysis revealed significant enrichment (FDR *q* < 0.05) in adhesion-related pathways, including *homophilic cell–cell adhesion via plasma-membrane adhesion molecules, cell–cell adhesion via plasma-membrane adhesion molecules*, and the related molecular function *calcium ion binding* (a hallmark of cadherin-mediated adhesion). These terms describe the junctional programs that preserve epithelial and endothelial barrier integrity. In aging, barrier function and junctional organization are known to decline, manifesting as increased permeability and altered intercellular communication, so the observed enrichment is biologically coherent with age-linked tissue changes and supports the plausibility of our edits acting through adhesion programs rather than core promoter CpG islands (see Figure 6).

Indeed, intervention CpGs were enriched in CpG islands and their shores but were depleted at promoter annotations, indicating a preference for intragenic/first-exon islands and distal regulatory contexts rather than canonical TSS cores. These two orthogonal lines of evidence strongly support the biological plausibility of our model. The Revive controller not only achieves its rejuvenation goal according to internal metrics but does so by prescribing edits to a concise set of loci that are demonstrably enriched for known aging biomarkers, located in key gene regulatory regions, and associated with the fundamental biological pathways that govern the aging process.

## 4 Conclusion

### Summary

Revive goes beyond epigenetic clocks to propose actionable CpG interventions that steer a methylome along a natural rejuvenation path. We learn a smooth, age-calibrated vector field in a stable PCA space and steer it with a convex, sparsity-aware controller. Integrated backward, the field defines a natural rejuvenation path; the controller then finds the smallest edit set that moves each sample toward it. On the EPIC–Italy hold-out, commanded rejuvenation translated linearly into realized age reduction (≈ 0.4 years per commanded year at low sparsity) with a clear sparsity–effect trade-off. Negative controls erased the effect, inferred cell-type composition remained stable, and targeted CpGs were enriched in island/shore and first-exon/body contexts and in adhesion-related pathways: signals consistent with known aging hallmarks. Crucially, Revive derives its supervisory signal from the methylome itself, obtaining labels directly from the features via age-conditioned flow learning, so it functions as a practical foundation model for epigenetic aging.

### Outlook

A natural next step is to train an aging foundation model conditioned not just on age and sex as we have done, but tissue, genotype, environment, and clinical covariates as well. Next, establishing rigorous and open benchmarks will be *sine qua non* for fair comparison and real progress. Finally, we encourage the community to build foundation models of epigenetic aging, leveraging the vast public troves of methylomes, metadata, sequences, and ages that are already available and naturally suited to large-scale self-supervision.

## A Appendix

### A.1 Algorithms

#### Algorithm 1

Preprocessing and Dynamics Model Training

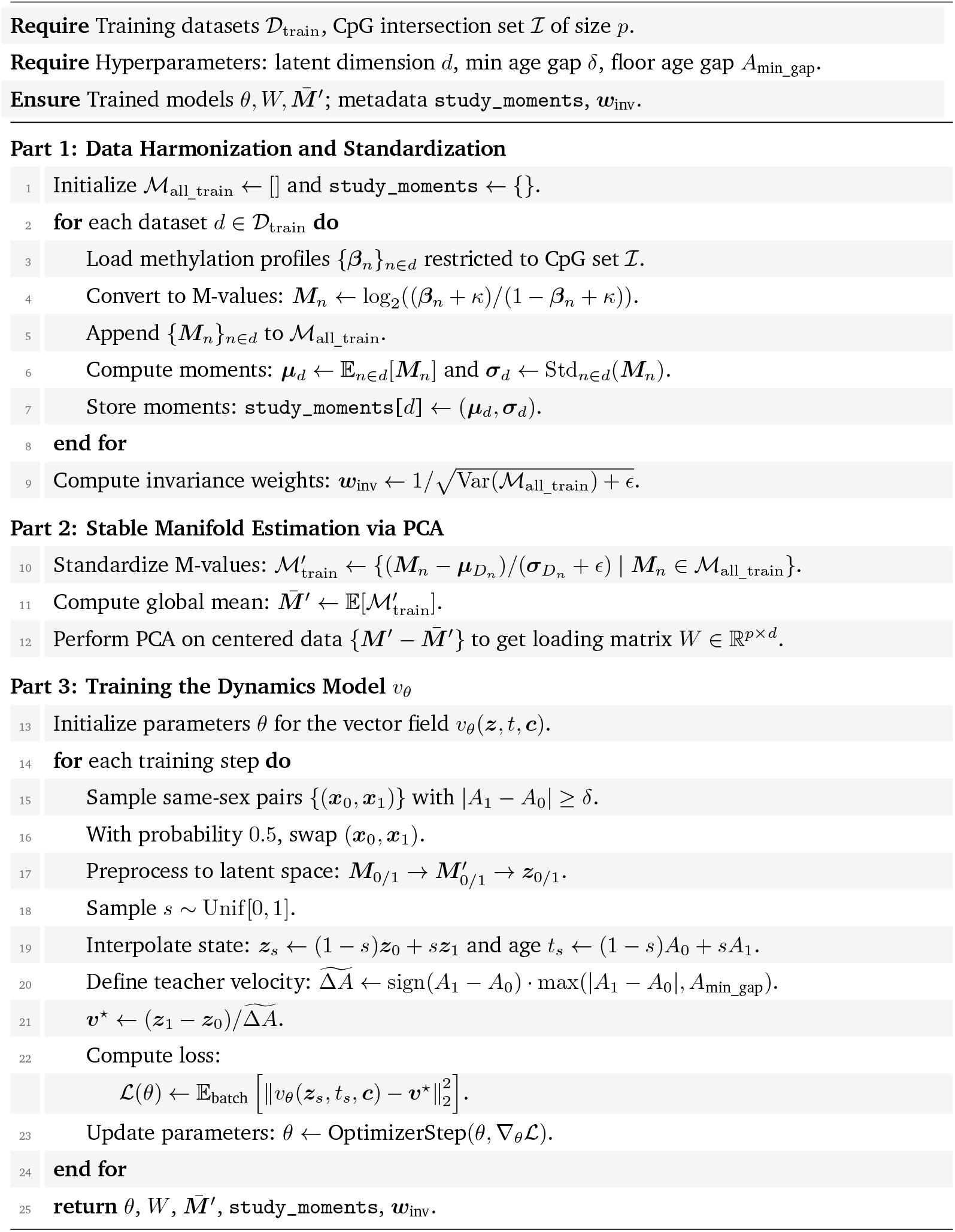

#### Algorithm 2

Minimal Intervention Generation via ADMM

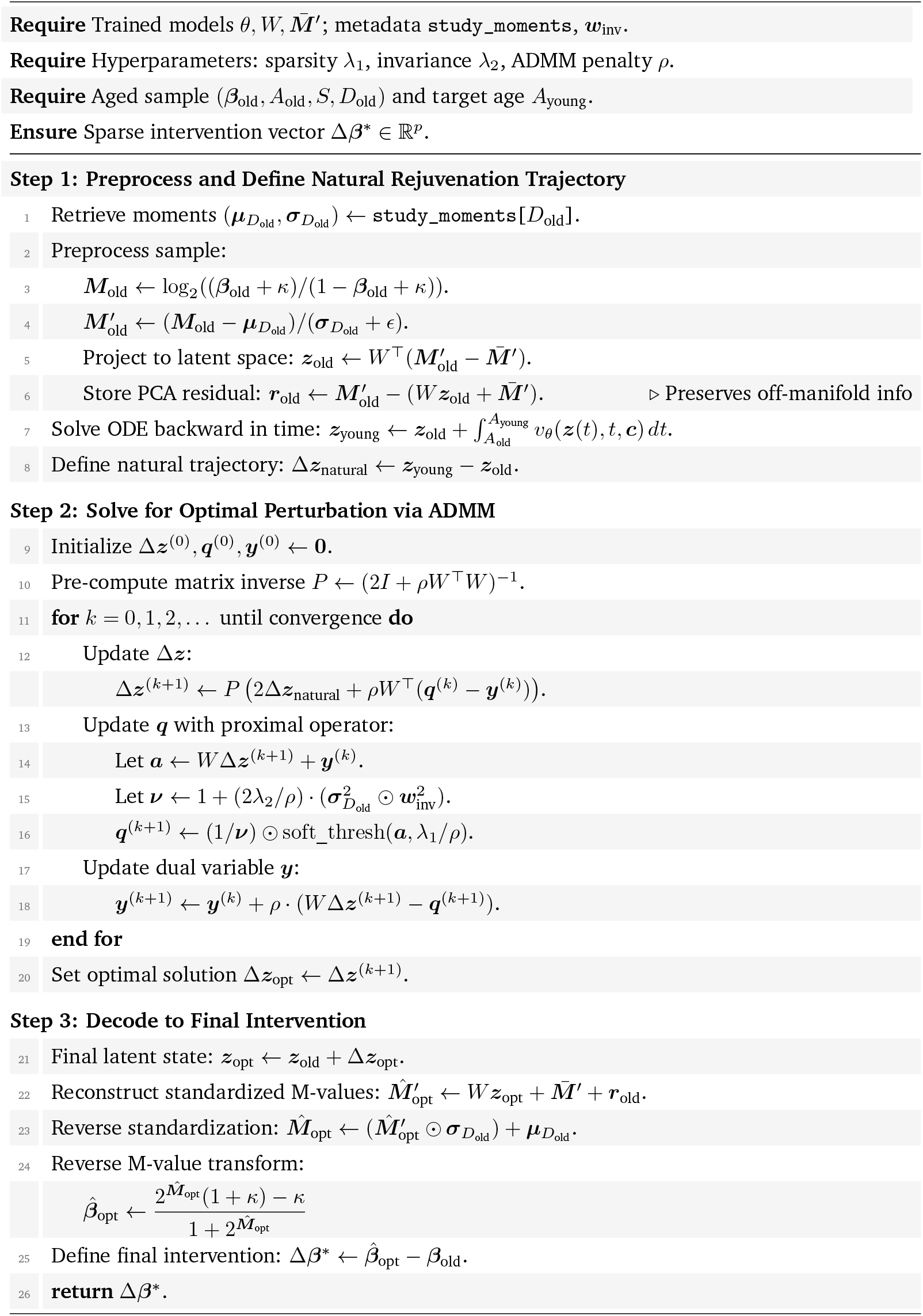

## Footnotes

a While an adaptive solver like RK45 could also be used, we use a fixed-step scheme for deterministic reproducibility.

